# The key features of SARS-CoV-2 leader and NSP1 required for viral escape of NSP1-mediated repression

**DOI:** 10.1101/2021.09.13.460054

**Authors:** Lucija Bujanic, Olga Shevchuk, Nicolai von Kügelgen, Katarzyna Ludwik, David Koppstein, Nadja Zerna, Albert Sickmann, Marina Chekulaeva

## Abstract

SARS-CoV-2, responsible for the ongoing global pandemic, must overcome a conundrum faced by all viruses. To achieve its own replication and spread, it simultaneously depends on and subverts cellular mechanisms. At the early stage of infection, SARS-CoV-2 expresses the viral nonstructural protein 1 (NSP1), which inhibits host translation by blocking the mRNA entry tunnel on the ribosome; this interferes with the binding of cellular mRNAs to the ribosome. Viral mRNAs, on the other hand, overcome this blockade. We show that NSP1 enhances expression of mRNAs containing the SARS-CoV-2 leader. The first stem-loop (SL1) in viral leader is both necessary and sufficient for this enhancement mechanism. Our analysis pinpoints specific residues within SL1 (three cytosine residues at the positions 15, 19 and 20) and another within NSP1 (R124) which are required for viral evasion, and thus might present promising drug targets. Additionally, we carried out analysis of a functional interactome of NSP1 using BioID and identified components of anti-viral defense pathways. Our analysis therefore suggests a mechanism by which NSP1 inhibits the expression of host genes while enhancing that of viral RNA. This analysis helps reconcile conflicting reports in the literature regarding the mechanisms by which the virus avoids NSP1 silencing.

## INTRODUCTION

SARS-CoV-2, responsible for the current global pandemic, manages to evade mechanisms of host immunity during infections to promote its own replication and spread. To achieve this, it simultaneously suppresses the translation of cellular proteins and promotes that of its own, although the two processes require the same basic machinery. Exposing the mechanisms by which it manages this would likely provide insights into crucial stages in the viral lifecycle that might be exploited in therapies.

SARS-CoV-2 belongs to the genus beta-coronavirus (1), which also includes a bat coronavirus (96% identity at the genome level with SARS-CoV-2), SARS-CoV-1 (or SARS-CoV), which caused SARS epidemic in 2003, and Middle East respiratory syndrome (MERS) CoV, responsible for an outbreak of respiratory disease in 2012 (reviewed in (2,3)). The genome of SARS-CoV-2 is a ~30 kb positive-stranded RNA with 5′-cap structure, 5′UTR (or leader), 3′UTR and polyA tail. It contains 10 protein-coding open reading frames (ORFs). Upon cell entry, the genomic RNA (gRNA) is translated into polyprotein, which is processed into 16 nonstructural proteins (NSPs). Subsequently, its gRNA serves as a template to generate a set of subgenomic mRNAs (sgRNAs) that encode other viral proteins. Curiously, all sgRNAs possess a common 5′ leader sequence, because the templates are switched from ORF-containing regions to the leader during sgRNA synthesis, by viral RNA-dependent RNA polymerase.

The first protein produced by coronaviruses upon infection is NSP1, encoded by ORF1a at the 5′ end of gRNA. NSP1 is an important virulence factor that plays a crucial role in its pathogenicity by helping the virus evade the host innate immune response (reviewed in (3)). In the related virus SARS-CoV-1, NSP1 inhibits immunity via two mechanisms: by repressing expression of host transcripts (4–9) and by preventing full induction of interferon (IFN) and decreasing STAT1 phosphorylation (10).

A number of studies have been devoted to determining how SARS-CoV-1 NSP1 represses host gene expression, but some of their conclusions seem contradictory (4–9). Two repression mechanisms have been reported: translational repression and mRNA degradation (6–8). Specific amino acid residues important for NSP1-mediated repression have been identified. NSP1 carrying [K164A; H165A] mutations is fully nonfunctional (7,9), and another [R124A; K125A] mutant lacks the mRNA destabilization function (7). The mechanism by which SARS-CoV-1 NSP1 achieves translational repression is not fully understood. Its effects are thought to relate to the general translation machinery, due to its co-sedimentation with the small ribosomal subunit (40S) and coimmunoprecipitation with the ribosomal protein S6 (6). Yet experiments with the separation of translation complexes on sucrose density gradients and toeprinting analyses produced conflicting data. While the former suggested that NSP1 inhibits recruitment of the large ribosomal subunit and formation of 80S initiation complex, the latter indicated that it rather affected the recruitment of the small (40S) ribosomal subunit and assembly of the 48S initiation complex (6). Additionally, some reports suggested that SARS-CoV-1 NSP1 affects only host mRNA, while SARS-CoV-1 mRNAs are protected from translational downregulation through interactions between virus-specific leader sequences with NSP1 (9). Other studies suggested that viral mRNAs are also translationally inhibited by NSP1 in SARS-CoV-1-infected cells, providing an overall picture that is confusing (7,11).

The COVID-19 pandemic has triggered intensive research into the mechanisms of NSP1 functions in SARS-CoV-2 (12–16), Biorxiv: https://doi.org/10.1101/2020.09.18.302901, https://doi.org/10.1101/2021.05.28.446204). Two cryo-EM studies showed that SARS-CoV-2 NSP1 binds to the ribosomal 40S subunit and blocks the mRNA entry tunnel (12,13). However, in order for the virus to propagate, viral translation has to proceed in the presence of NSP1. Here, too, attempts to resolve the underlying mechanisms have produced conflicting results. As with studies on SARS-CoV-1 NSP1, some work has reported that SARS-CoV-2 NSP1 represses both host mRNAs and mRNAs with viral leader (13); other studies found that viral reporters escape repression by NSP1 (15,16), Biorxiv: https://doi.org/10.1101/2021.05.28.446204). When the evasion of viral reporters from NSP1-mediated repression has been reported, authors have disagreed about the viral elements that are required. Shi et al. (Biorxiv: https://doi.org/10.1101/2020.09.18.302901) reported that multiple elements in viral leader make contributions, while Tidu et al. (15) and Banerjee et al. (16) argue that a specific stem-loop structure in the viral leader suffices.

Here we use a combination of reporter assays, mutagenesis and mass spectrometry to dissect the mechanisms of SARS-CoV-2 NSP1 function and provide insights into how the virus evades NSP1 silencing. We show that SARS-CoV-2 NSP1 both downregulates global protein production and fosters the expression of viral reporters. We find that the stem-loop 1 (SL1) in viral leader is both necessary and sufficient for upregulation of viral reporters. We map three specific cytosine residues (3C) within SL1 and an arginine residue at the position 124 in NSP1 which are absolutely required for viral evasion. Mutation of any of these four residues, alone or in combination, is sufficient to make the viral reporter susceptible to NSP1 repression. Moreover, we use BioID (17) to determine the functional interactome of SARS-CoV-2 NSP1, identifying multiple components of the anti-viral defense system.

## MATERIALS AND METHODS

### Cell culture, transfections, and luciferase assay

Human HEK293T cells were grown in Dulbecco’s modified Eagle’s medium with GlutaMAX™ supplement (DMEM+ GlutaMAX, GIBCO) with 10% FBS. Transfections were done in 96-well plates with polyethylenimine (PEI) using a 1:3 ratio of DNA:PEI. In reporter experiments, HEK293T cells were transfected with 1-2 ng RL, 15 ng FL, and indicated amounts of NSP1-encoding constructs per well of a 96-well plate. Total amount of transfected DNA was topped up to 50 ng per well of 96-well plate with the vector. Cells were lysed 24 hr post transfection. Luciferase activities were measured with a homemade luciferase reporter assay system as described earlier (18).

### DNA constructs

3xflag-SARS-CoV-2 NSP1-encoding plasmid and pEBG-3xflag, used as a vector, have been described previously (19). Analogous plasmid expressing SARS-CoV-1 NSP1 was generated using a similar strategy: CDS of NSP1 was PCR amplified, using SARS-CoV-1 cDNA as a template, and cloned between SbfI and NotI sites of pEBG-3xflag. R124A, [R124A; K125A], and [K164A; H165A] mutations were introduced in SARS-CoV-2 and SARS-CoV-1 NSP1 CDS by site-directed mutagenesis. RL reporter is similar to previously described RL plasmid (20), but lacks the last 8 nt in the CMV promoter, which were removed by site-directed mutagenesis. RL served as a backbone for cloning of Renilla reporters carrying SARS-CoV-2 leader, as well as its deletion and point mutants. SARS-CoV-2 leader (attaaaggtttataccttcccaggtaacaaaccaaccaactttcgatctcttgtagatctgttctctaaacgaacaaactaaaatgtctgataatggacccca) was generated by oligo annealing and cloned between SacI and NheI sites of RL, to produce CoV-2-RL. For CoV-2-ΔSL1-RL, leader lacking the first 33 nt (aaccaactttcgatctcttgtagatctgttctctaaacgaacaaactaaa) was cloned upstream of RL. SL1-RL (or CoV-2-SL-RL) and CoV-1-SL1-RL contains the first 33 nt of SARS-CoV-2 (attaaaggtttataccttcccaggtaacaaacc) or the first 31 nt of SARS-CoV-2 leader (atattaggtttttacctacccaggaaaagcc), correspondingly. Mutations of SL1, indicated in the figures, were introduced into the oligos used for cloning. To clone gCoV-2-flag-NSP1 plasmids, CDS of NSP1 and its point mutants were PCR amplified and cloned between NheI and NotI sites of RL, to substitute RL CDS. At the next step, SARS-CoV-2 genomic leader was PCR amplified using SARS-CoV-2 cDNA as a template (5’ end: attaaaggtttataccttcccagg; 3’ end: cttacctttcggtcacacccggac) and cloned it between SacI and NheI of NSP1 plasmids. For BioID constructs, BioID CDS was PCR amplified from pcDNA 3.1-BioID (17) and cloned between BstXI and NotI of pEBG-sic (21), to produce pEBG-BioID. pEBG-BioID was used as a mock control in BioID experiments and as a backbone for cloning NSP1-myc-BioID constructs. For that, CDS of NSP1 or its point mutants were PCR amplified and cloned between NheI and SmaI sites of pEBG-BioID.

### Western blotting and BioID

For western blotting, 20 μl of total cell lysate from reporter assay was separated on a 10 % SDS-PAGE, and proteins were transferred to the PVDF membrane. The membrane was probed with the following primary antibodies: mouse anti-flag antibody 1:2000 (F1804 Sigma), mouse anti-beta-actin 1:5000 (A2228 Sigma).

For BioID experiments, HEK293T cells were transfected with constructs encoding NSP1-BioID, NSP1-KH164AA-BioID, NSP1-RK124AA-BioID or BioID alone. Transfections were done in quadruplicates, using 10 μg of plasmid and 30 μg of polyethylenimine (PEI) per 10 cm dish with 3×10^6^ cells plated a day before transfection. Cell culture medium was supplemented with 50 μM biotin. Cells were lyzed 24 hr post-transfection and BioID was performed as previously described (22). In short cells were lyzed in 8M urea, 50mM Tris-HCl pH 7.4, 1x protease inhibitors Complete EDTA-free, 1mM DTT. Lysates were supplemented with Triton-X100 to final concentration 1 %, sonicated, diluted 5-fold with lysis buffer and clarified. Biotinylated proteins were isolated by incubation with 100 μl of streptavidin dynabeads (Thermo 65001) at 4°C with rotation overnight. Proteins were eluted 2x with 25 μl of elution buffer (5% SDS, 50 mM Tris-HCl pH 7.4) and used for mass spectrometry analysis.

### Mass spectrometry: In solution digestion and LC-MS/MS analysis

The BioID IP eluates (10 μL each) were diluted in ultra-pure water whereas cell lysates each corresponding to 20 μg of protein were diluted in 50 mM ammonium bicarbonate (NH_4_HCO_3_) buffer, pH 7.8 to a final volume of 100 μL. All samples were reduced with 10 mM DTT at 56°C for 30 min, and subsequently alkylated with 30 mM IAA at room temperature for 30 min in the dark. Next, the samples were subjected to ethanol (EtOH) precipitation followed by in-solution protein digestion. Briefly, each sample was diluted 10-fold with ice-cold EtOH in 1:10 (v/v) ratio, vortexed and incubated at −40°C for 60 min followed by centrifugation in a pre-cooled (4°C) centrifuge at 20,000 g for 30 min. The obtained pellet was washed with 100 μL of ice-cold acetone, briefly vortexed and centrifuged as mentioned above for 5 min. The supernatant was discarded, the pellet was dried under laminar flow hood and re-solubilized in 60 μL of digestion buffer comprising: 0.2 M GuHCl, 2 mM CaCl_2_, 50 mM NH_4_HCO_3_, pH 7.8. 100 ng of Trypsin Gold and 1 μg of Trypsin sequenced grade (both Promega) were added to BioID eluates and total cell lysates, respectively and subjected to proteolysis at 37°C for 16 h. Lastly, all samples were acidified with 10% TFA to pH < 3.0 and an aliquot of each digest i.e. 10% of BioID and 5% of total cell lysate was quality controlled as described previously (23).

For LC-MS/MS analysis, 30% of BioID eluates and 10% of total cell lysates digests were analyzed using an Ultimate 3000 nano RSLC system coupled to Orbitrap Lumos (both Thermo Scientific). Peptides were pre-concentrated on a 100 μm x 2 cm C18 trapping column for 5 min using 0.1% TFA with a flow rate of 20 μL/min followed by separation on a 75 μm x 50 cm C18 main column (both Acclaim Pepmap nanoviper, Thermo Scientific) with a 60 min (BioID samples) or 120 min (cell lysate samples) LC gradient ranging from 3-35% of B (84% ACN in 0.1% FA) at a flow rate of 250 nL/min. The Orbitrap Lumos was operated in data-dependent acquisition mode and MS survey scans were acquired from m/z 300 to 1500 at a resolution of 120000 using the polysiloxane ion at m/z 445.12002 as lock mass (24). For MS1 scans, the automatic gain control (AGC) target value was set to 2 x 10^5^ with a maximum injection time (IT) of 50 ms. MS2 spectra were acquired in the linear ion trap (rapid scan mode) after higher-energy collisional dissociation with a normalized collision energy of 30% and an AGC target value of 2 x 10^3^ and a maximum IT of 300 ms, by utilizing a maximal duty cycle of 3 s, prioritizing the most intense ions and injecting ions for all available parallelizable time. Selected precursor ions were isolated using quadrupole with a 1.2 m/z window taking into account a dynamic exclusion of 30 s.

For data analysis, all MS raw data were processed with Proteome Discoverer software 2.3.0.523 (Thermo Scientific, Germany) and searched in a target/decoy fashion against a concatenated version of the human Uniprot database (downloaded on on November 2019, 20300 target sequences); NSP1 from SARS-CoV-2 and BioID SEQUEST-HT algorithm. The search parameters were: precursor and fragment ion tolerances of 10 ppm and 0.5 Da for MS and MS/MS, respectively. Trypsin was set as enzyme with a maximum of 2 missed cleavages.

Carbamidomethylation of Cys as fixed modification and oxidation of Met was selected as dynamic modification. The false discovery rate was set to 0.01 for both peptide and protein identifications using Percolator. A Label-free quantification (LFQ) analysis was performed with four replicates for each condition for whole proteome analysis and for pull down experiment. Proteins identified with ≥ 2 unique peptides were used for differential expression analysis. Enrichment (log2 fold change) of proteins between pulldown fractions or lysate samples was calculated using a generalised linear model (R limma package, (25)) on imputed log2-transformed LFQ values. P-values were adjusted for multiple testing using the FDR method.

## RESULTS

### Stem-loop 1 (SL1) is both necessary and sufficient for NSP1-mediated upregulation of viral RNA expression

To recapitulate SARS-CoV-2 NSP1-mediated repression in HEK293T cells, we set up a luciferase reporter assay. We co-expressed Renilla luciferase mRNA (RL) with NSP1-encoding constructs (**Figure 1A**). As negative controls, we used an empty vector and a NSP1 KH164AA [K164A; H165A] mutant reported as nonfunctional due to a disruption of interactions with the ribosome (12–14). As expected, WT NSP1, but not its KH164AA mutant, efficiently repressed luciferase expression in a dose-dependent manner (4- to 30-fold, **Figure 1B**). Because SARS-CoV-1 RK124AA [R124A; K125A] NSP1 mutant has been reported to disrupt the NSP1-mediated destabilization of mRNAs (7), we also included RK124AA and R124A mutants in the analysis. Both mutants were able to repress mRNA expression, although to a lesser degree than WT (1.5- to 7-fold, **Figure 1B**). Similar behaviour was observed for SARS-CoV-1 NSP1 and its mutants (**Figure 1C**). Consistent with a role of NSP1 in global translational repression, the expression levels of WT NSP1 and its mutants anti-correlated with their strength as translational repressors: WT NSP1 (from both SARS-CoV-2 and 1) was expressed the lowest, R124A and RK124AA mutants had intermediate expression levels, and KH164AA mutant was expressed the highest (**Figure 1B-C**).

**Figure 1.**
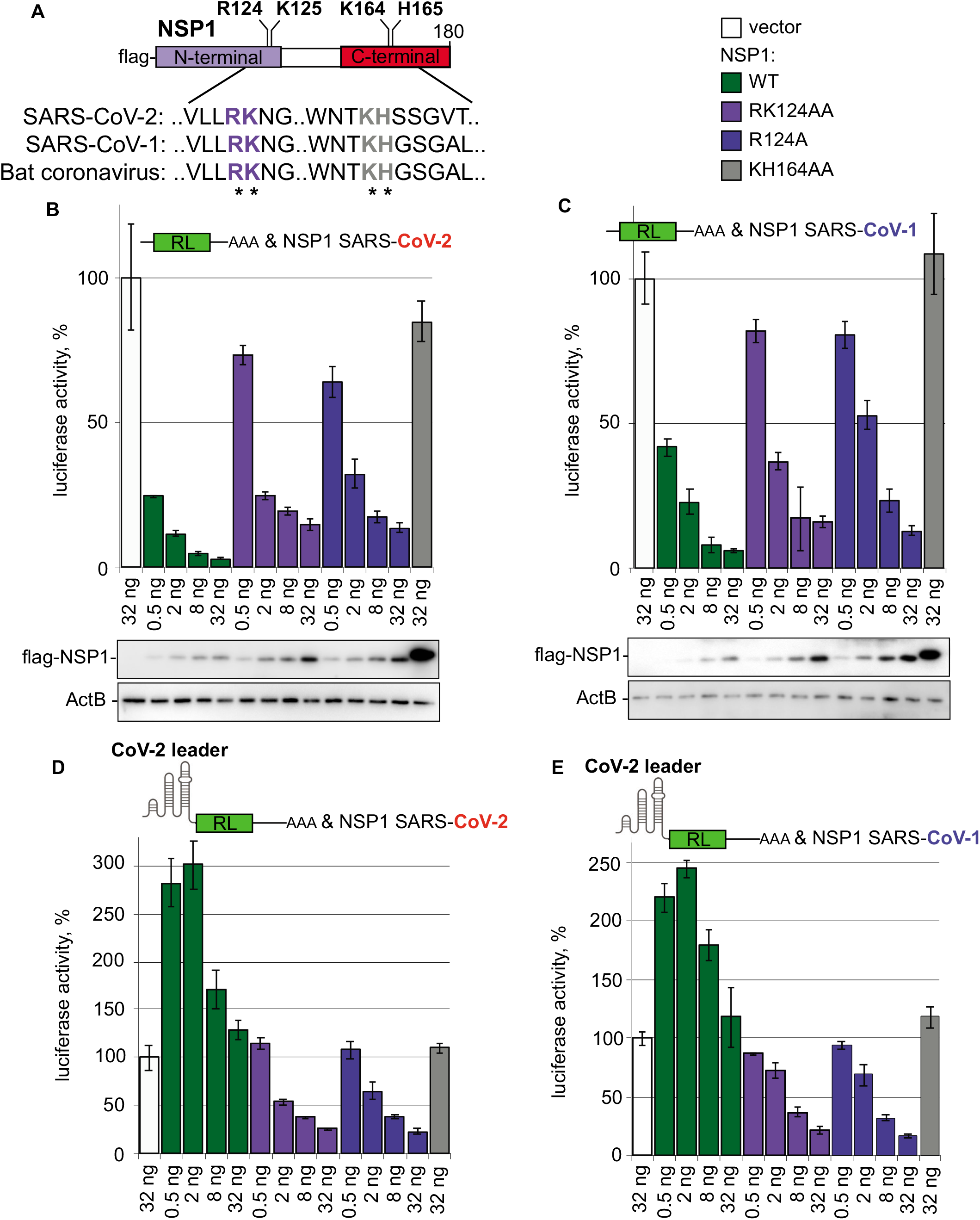
SARS-COV-2 5’UTR enables virus to escape repression by NSP1 protein, but not its R124A mutant. (**A**) Schematic representation of flag-tagged NSP1 protein and analyzed mutants (R124A, RK124AA, KH164AA), with partial alignment of mutated regions. The numbers correspond to the amino acid positions. (**B**) Repression of RL mRNA by SARS-CoV-2 NSP1 and its point mutants. HEK293T cells were co-transfected with Renilla luciferase (RL) plasmid and increasing amounts of plasmids, encoding flag-tagged SARS-CoV-2 NSP1 or indicated NSP1 point mutants. As negative control, vector encoding flag alone instead of flag-NSP1 plasmid was used. Open bars: vector, green: WT NSP1, purple: RK124AA NSP1, blue: R124A NSP1, grey: KH164AA NSP1. Values are presented as a percentage of luciferase produced in the presence of vector. Values represent means +/-SD from at least 3 experiments. Expression of flag-NSP1 fusion protein and its point mutants was estimated by western blotting with anti-falg antibodies and shown below the reporter assay. ActB was used a loading control. (**C**) Repression of RL mRNA by SARS-CoV-1 NSP1 and its point mutants. The experiment was performed as in (B), but SARS-CoV-1 NSP1 instead of SARS-CoV-2 NSP1 was used. (**D-E**) Reporter bearing SARS-CoV-2 leader (CoV-2-RL) escapes repression by WT NSP1, but is repressed by NSP1 R124A mutant. Panel (**D**) shows the effects of NSP1 from SARS-CoV-2 and panel (**E**) – NSP1 from SARS-CoV-1. 5′UTR corresponds to the subgenomic RNA encoding nucleocapsid protein (ORF9).

The mRNAs of SARS-CoV-1 (9) and SARS-CoV-2 (15,16) Biorxiv: https://doi.org/10.1101/2020.09.18.302901, https://doi.org/10.1101/2021.05.28.446204) have been reported to escape NSP1-mediated repression through interactions with NSP1 itself. To recapitulate this process, we added SARS-CoV-2 leader to our Renilla luciferase reporter (CoV-2-RL). Under this condition, the expression of CoV-2-RL mRNA was not repressed (**Figure 1D**). Moreover, low doses of WT NSP1 stimulated the expression of CoV-2-RL at a level of 2.5-to-3-fold. In SARS-CoV-1, the R124A NSP1 mutant has been reported to be defective in its binding to the viral leader (9). Therefore, we tested whether SARS-CoV-2 RK124AA and R124A mutants were still able to repress CoV-2-RL. Strikingly, both RK124AA and R124A mutants continued to repress CoV-2-RL. We observed similar behaviour for SARS-CoV-1 NSP1 and its mutants (**Figure 1E**).

We next decided to determine whether a specific region of the SARS-CoV-2 leader is responsible for alleviation of NSP1 silencing. To achieve this, we performed a deletion analysis of SARS-COV-2 leader in reporter assay. Given earlier reports on the role of stem-loop 1 (SL1) in the expression of SARS-CoV-1 (9) and SARS-CoV-2 (15,16) Biorxiv: https://doi.org/10.1101/2020.09.18.302901), we generated reporters in which SL1 is deleted (CoV-2 ΔSL1-RL), or which contain only the SL1 region (CoV-2 SL1-RL, CoV-1 SL1-RL). This analysis showed that SL1 is both necessary and sufficient to escape NSP1-mediated repression (**Figure 2A**). Consistently, when we generated NSP1-encoding constructs carrying native SARS-CoV-2 leader, we observed that WT NSP1 was expressed at higher levels than its point mutants (**Figure 2B**). These results suggest that viral-encoded NSP1 enhances its own expression and that of other viral proteins, while inhibiting the expression of host mRNAs.

**Figure 2.**
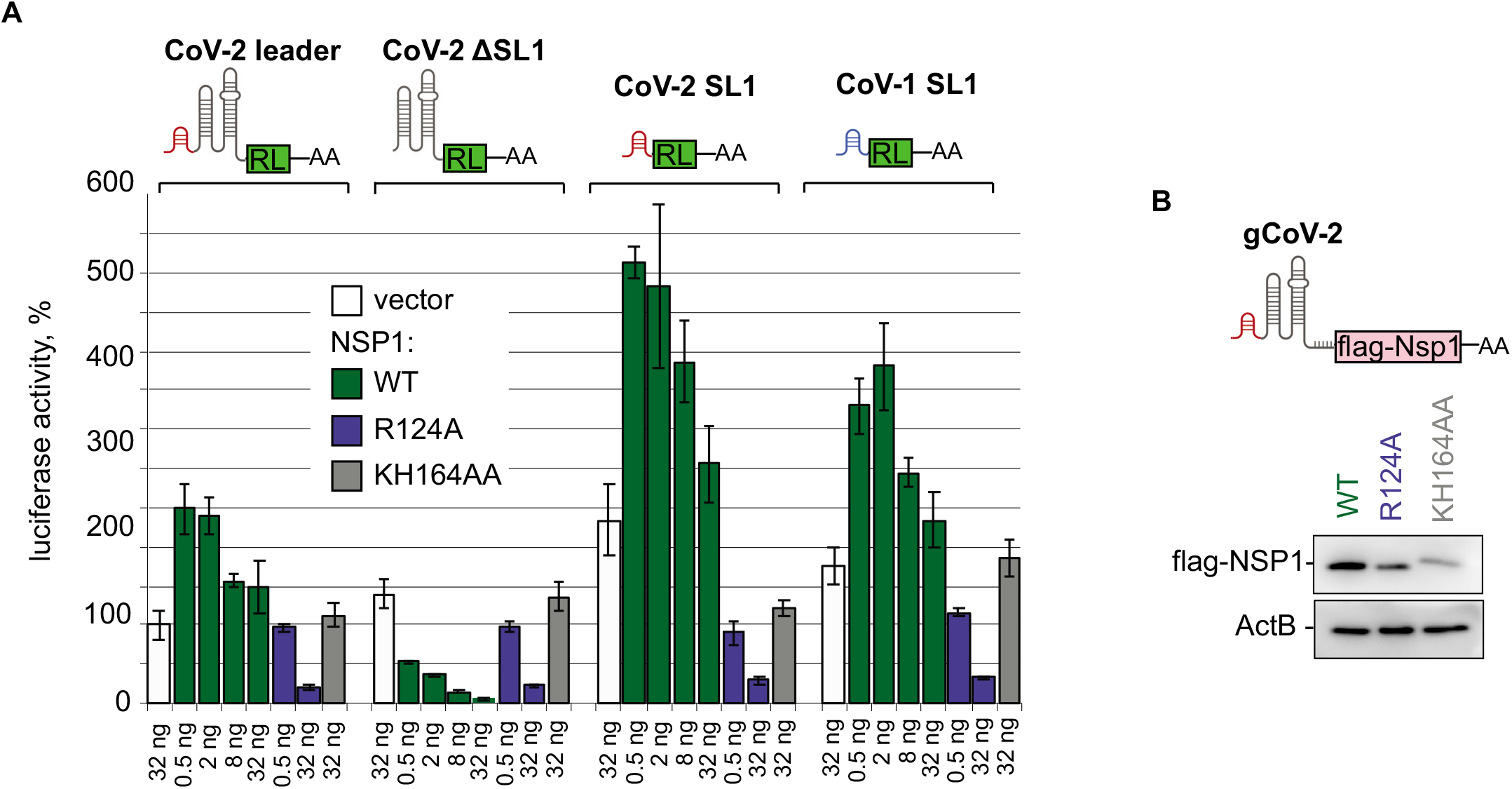
SARS-COV-2 leader alleviates NSP1-mediated silencing via its stem-loop 1 (SL1) structure and enhances NSP1 expression. (**A**) SL1 in SARS-COV-2 leader is both necessary and sufficient to escape NSP1-mediated repression. HEK293T cells were co-transfected with the indicated SARS-COV-2 leader reporters and SARS-CoV-2 NSP1-encoding constructs, either WT or indicated mutants. CoV2-RL reporter contains full-length leader of subgenomic SARS-CoV-2 RNA (ORF9) upstream of RL coding sequence, CoV-2-ΔSL1-RL lacks stem-loop 1 (SL1), and CoV2-SL1-RL and CoV-1-SL1-RL carry SL1 alone, originating from SARS-CoV-2 and SARS-CoV-1, correspondingly. The experiment was performed and data presented as in Figure 1B. (**B**) NSP1 encoded by mRNA carrying viral leader enhances its own expression. Constructs encoding NSP1 or its point mutants and carrying genomic leader of SARS-CoV2 (gCoV-2-flag-NSP1) were transfected in HEK293T cells, and cell lysates were analyzed by western blotting with anti-flag antibody. Beta-actin was used as a loading control.

### Three cytosine residues in SL1 are necessary for its derepressor function

In a next step, we mapped the residues within SL1 which are required for its function as a derepressor. To this end, we carried out extensive mutagenesis of SL1 and tested how specific mutations affected the expression of the SL1-RL reporter in the presence of NSP1. SL1 is highly conserved between SARS-CoV-2, bat CoV and SARS-CoV-1, while MERS-CoV SL1 shows less conservation (**Figure 3A**). SARS-CoV-2 SL1 consists of two 10 bp-long double helices (stem 1a and 1b), with a bulge in between, and a 4 nt-long loop. Given the conservation of the loop sequence (U/ACCC), we first mutated the residues within it and tested how these mutations affected the ability of SL1 to escape NSP1-mediated repression in the luciferase reporter assay. The 18U>A (*i.e*. at the position 18 of SL1 a U was changed to an A) and 21C>G mutants were functional. But mutations of 19C and 20C, individually or in combination (19C>G, 20C>G, [19C>G; 20C>G], [19C>G; 20C>G; 21C>G], [18U>A; 19C>G; 20C>G; 21C>G]), disrupted the derepressor function of SL1 (**Figure 3B**).

**Figure 3.**
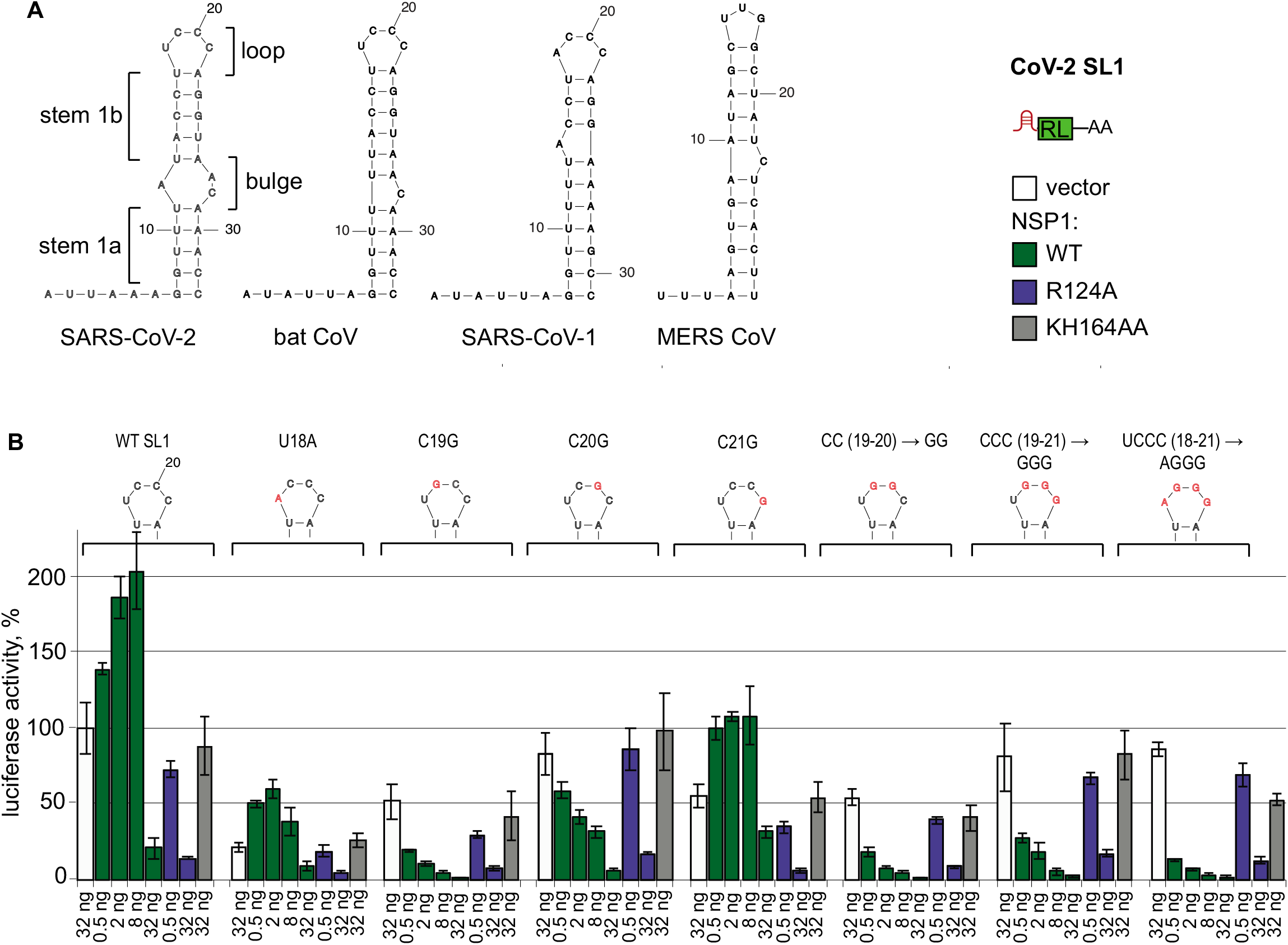
Positions C19 and C20 in the loop of SARS-CoV-2 SL1 are required to escape NSP1-mediated repression. (**A**) Structures of SARS-CoV-2, bat CoV, SARS-CoV-1, and MERS-CoV SL1, predicted by mfold. The elements of SL1 are labelled: loop, two sections of stem 1 (stem1a and stem 1b), separated by bulge. (**B**) Mutagenesis of the loop shows the requirement of C19 and C20 for the derepressor function of SL1 in reporter assay. HEK293T cells were co-transfected with SL1-RL reporter or indicated SL1 mutants and the constructs encoding SARS-CoV-2 NSP1, either WT or indicated mutants. The introduced mutations (18U>A, 19C>G, 20C>G, 21C>G, [19C>G; 20C>G], [19C>G; 20C>G; 21C>G], [18U>A; 19C>G; 20C>G; 21C>G]) and resulting sequences of the loop are shown above the plots. The mutated residues are shown in red. The results are presented as in Figure 1B.

We next tested whether features of the stem such as its length or the presence of the bulge affect the function of SL1. We found that either extending or shortening the stem by 5 bp preserved part of the SL1 activity. NSP1 was not able to repress these reporters, although the overall efficiency of the expression of mutated SL1-RL and the degree of upregulation by NSP1 were on average lower than for WT SL1-RL (**Figure 4A**). Similar results were observed for SL1 without the bulge (**Figure 4A**).

**Figure 4.**
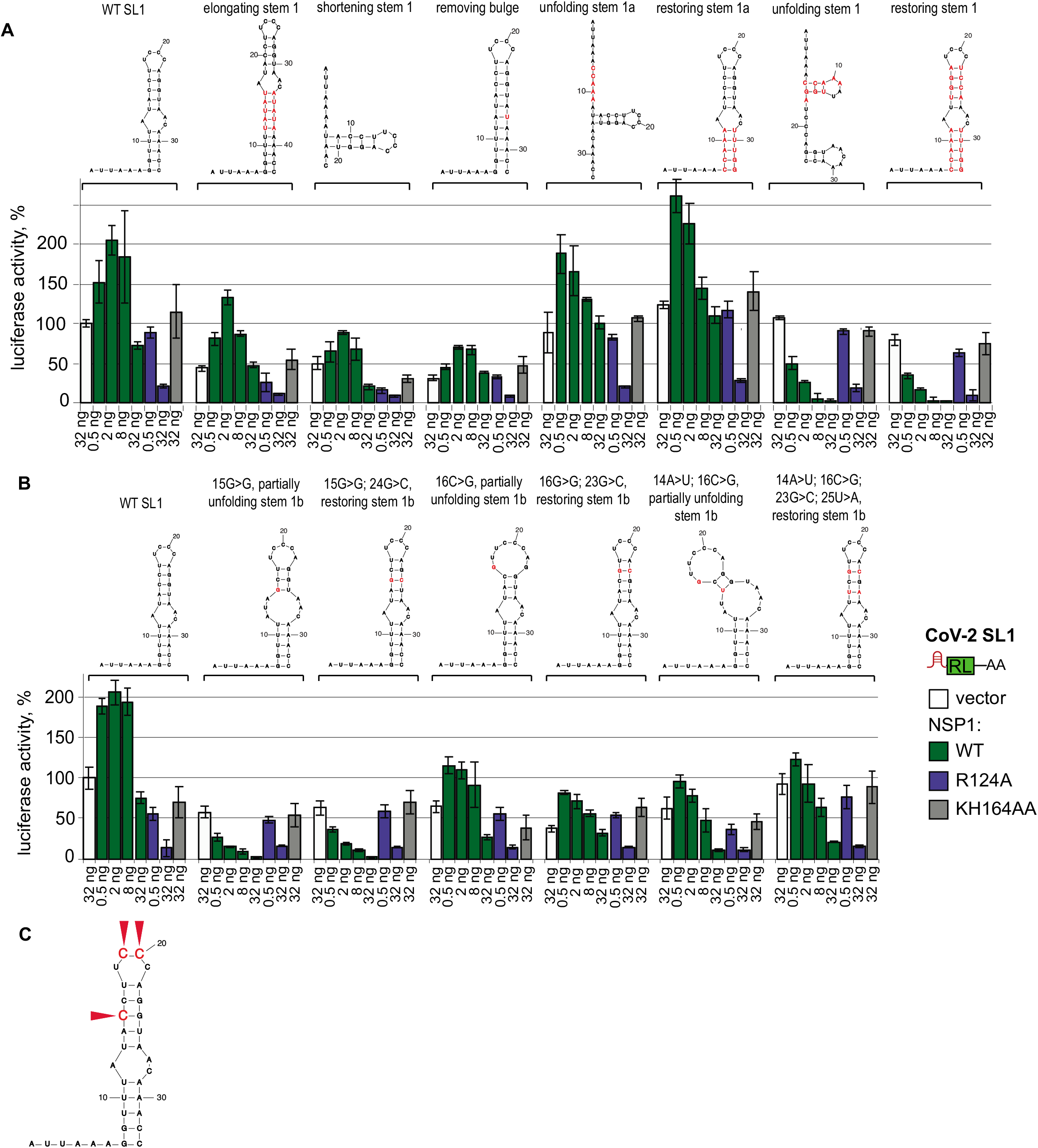
Position C15 in the stem of SARS-CoV-2 SL1 is necessary to escape NSP1-mediated repression. (**A**) Sequence of stem 1b (see Figure 2A) is important for derepressor function of SL1. The role of mutations, disrupting folding or modifying parameters of the stem and bulge in SL1, was tested in NSP1-reporter assay, as described in Figure 3B. Mutations and resulting predicted structures (mfold) are shown above the plots, with mutated residues in red: 7_11del GGUUUinsCCAAA (i.e replacement of nucleotides 7 to 11 (GGUUU) by CCAAA) unfolding stem 1a, [17_11delGGUUUinsCCAAA; 29_33delAAACCinsUUUGG] restoring stem 1a, [7_11del GGUUUinsCCAAA; 14_17delACCUinsUGGA] unfolding the stem 1, [7_11del GGUUUinsCCAAA; 14_17delACCUinsUGGA; 22_25delAGGUinsUCCA; 29_33delAAACCinsUUUGG] restoring stem 1, [12_16insUAUAU; 34_38AUAUA] elongating stem 1 by 5 bp, [7_11delCCAAA; 29-33del UUUGG] shortening stem 1 by 5 bp, 27_28delACinsU removing bulge. (**B**) Position 15C in stem 1b is required for SL1 function. Point mutations unfolding stem 1b, as well as compensatory mutations restoring folding, were introduced into SL1-RL reporter and tested in Nsp1-mediated repression assay, as described in (A). Mutations introduced in stem 1b: 15G>G unfolding; [15G>G; 24G>C] restoring folding; 16C>G unfolding; [16C>G; 23G>C] restoring folding; [14A>U; 16C>G] unfolding; [14A>U; 16C>G; 23G>C; 25U>A] restoring folding. (**C**) Three cytosine residues in SARS-CoV-2 SL1, one in stem 1b and two in the loop, are crucial to escape NSP1-mediate repression. Functional residues are marked with red arrows. The data are based on reporter assays shown in Figure 3B, 4A&B.

Our next question was whether the stem simply functions as a secondary structure that is required to present the loop in the right orientation, or whether the stem’s sequence is also functionally important. To test this, first we mutated one of the strands to unfold the stem, and then introduced compensatory mutations in the second strand to restore the secondary structure. Unfolding the first part of the stem, located prior to the bulge (**Figure 4A**, unfolding stem 1a) preserved most SL1 activity, consistent with our results from the shortening experiments. However, unfolding the entire stem fully abrogated the derepression activity of SL1 (**Figure 4A**, unfolding stem 1). Introducing compensatory mutations that restored the stem, did not bring back its activity. These results suggest that the structure of the stem *per se* is not sufficient for the derepressor function of SL1; instead, the sequence of the stem contributes to its function.

To explore which residues in the stem are important, we introduced point mutations. Because mutations of stem 1a were tolerated (**Figure 4A**), we mutated individual residues in stem 1b, adjacent to the loop. While mutations 16C>G and [14A>T; 16C>G] preserved much of the function, a mutation at position 15 fully abrogated SL1 function (15C>G, **Figure 4B**). Importantly, restoring the complementarity of the strands by introducing a compensatory mutation in the second strand of stem 1b did not restore stem activity ([15G>G; 24G>C], **Figure 4B**). This suggests that the specific sequence at this position is required for SL1 function.

To summarize, our analyses (**Figures 3** and **4)** identified three cytosines in SL1 which are required to provide for high expression of viral reporter: 15C, 19C and 20C (red arrows, **Figure 4C**).

### NSP1 interacts with ribosomal proteins, mRNA export and anti-viral defense components

To characterize the functional interactome of NSP1, we applied BioID (17), which outperforms pulldown assays in catching transient interactors. This assay relies on fusing the protein of interest with a promiscuous biotin ligase (birA*, **Figure 5A**). The ligase biotinylates any proteins in its close proximity (~ 10nm). These biotinylated proteins are subsequently purified on streptavidin beads and analyzed using mass spectrometry. We used this assay to identify the interactome of SARS-CoV-2 NSP1. As negative controls, we expressed biotin ligase alone (mock) or NSP1 mutants, KH164AA and RK124AA.

**Figure 5.**
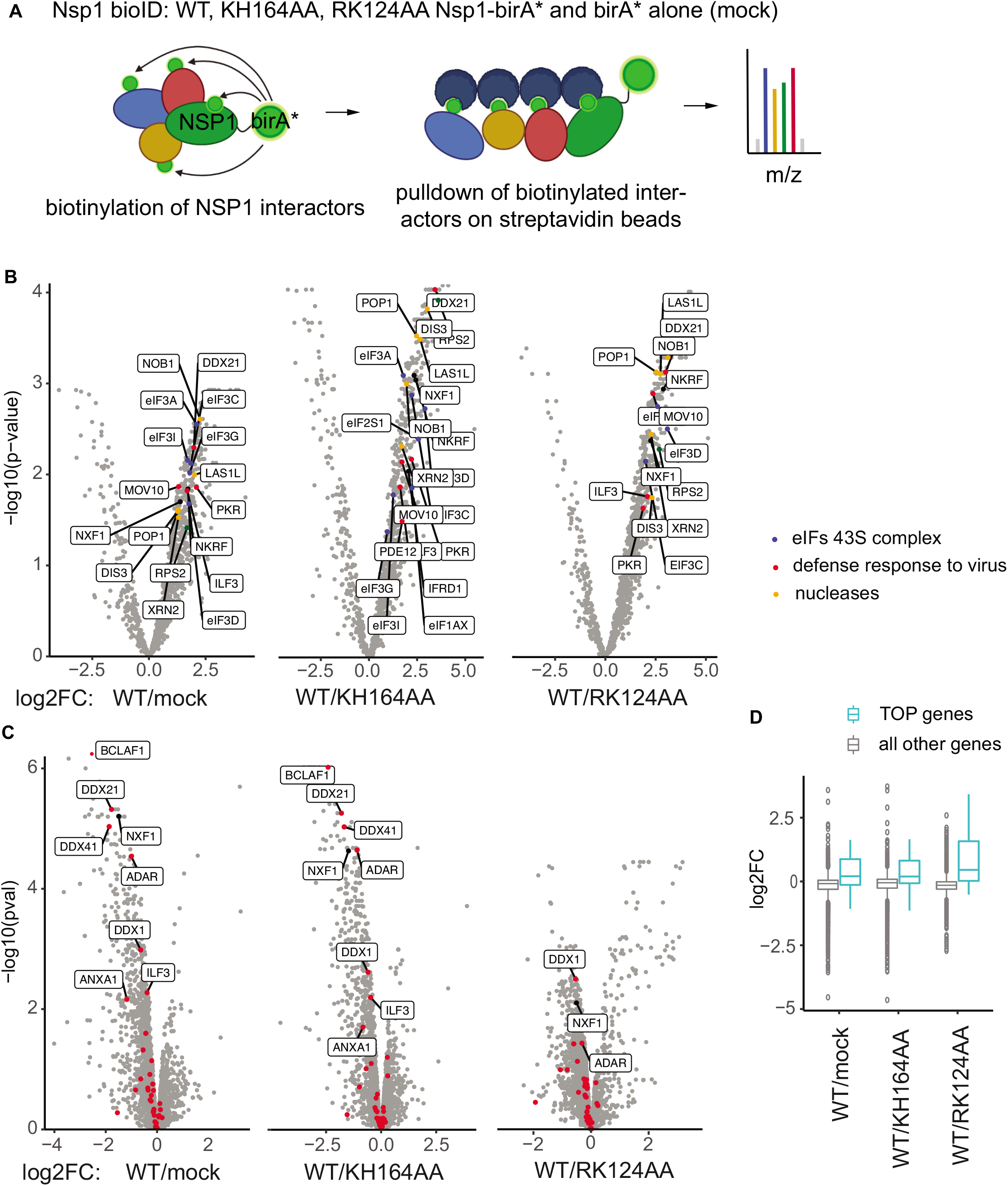
Nsp1 interacts with ribosomal proteins, ribosome biogenesis factors and viral response factors. (**A**) Scheme for NSP1 BioID. NSP1 is fused with a promiscuous biotin ligase (birA), leading to labelling of proteins within a ~ 10nm radius. Labelled proteins are isolated on streptavidin beads and quantified by LC-MS/MS. Unfunctional mutants of NSP1 and biotin ligase alone (mock) serve as negative controls. (**B**) Volcano plots showing proteins enriched in SARS-CoV-2 NSP1-BioID. -log10 p-values (Y) are plotted against log2 fold change of LFQ (label free quantification) values between NSP1-BioID and the indicated NSP1 mutant or BioID alone (mock) (X). Specific ribosomal protein (green), nuclear export factor 1 (NXF1, black), viral defense factors (referred to in the text and assigned to the GO category “defense response to virus’, red) and nucleases (orange) are shown. (**C**) Volcano plots showing changes in total proteome of 293T cells upon expression of SARS-CoV-2 NSP1. -log10 p-values (Y) are plotted against log2 fold change of LFQ (label free quantification) values between NSP1 and the indicated NSP1 mutant or tag alone (mock) (X). (**D**) Boxplot showing changes in expression of proteins encoded by TOP genes (cyan) and all other genes (grey) upon expression of WT NSP1, compared with mock (WT/mock), KH164AA NSP1 mutant (WT/KH164AA) and RK124AA mutant (WT/RK124AA).

NSP1 has been reported to inhibit host translation by inserting its C-terminal domain into the mRNA entry tunnel on the ribosomal 40S subunit (12–14). Consistent with these data, we detected interaction of WT NSP1 with PRS2/uS5 ribosomal protein situated near the mRNA entry tunnel (**Figure 5B**, NSP1/mock). Other ribosomal proteins, located within the radius of birA* activity (~ 10nm) from mRNA entry tunnel, were also detected among interactors. Other components of the 43S pre-initiation complex - initiation factors eIF1A, eIF2 and eIF3 - were also detected (**Figure 5B**, blue). Importantly, the KH164AA NSP1 mutant, reported to be defective in interaction with the ribosome (12–14), failed to interact with ribosomal proteins and initiation factors (**Figure 5B**, WT/KH164AA). RK124AA NSP1 mutant was able to repress mRNA, although not as efficiently as WT NSP1 (**Figure 1A, B**). Consistently, RK124AA mutant interacted with ribosomal proteins and initiation factors weaker than WT NSP1 (**Figure 5B**, WT/RK124AA).

Moreover, we found an enrichment of the nuclear export factor NXF1 in NSP1 BioID (**Figure 5B**), in line with the recent data showing that SARS-CoV-2 disrupts mRNA export from the nucleus (26,27). Additionally, the levels of NXF1 were downregulated in cells expressing WT NSP1, when compared with mock or KH164AA (**Figure 5C**). Besides changes in NXF1 levels, we observed that WT NSP1 upregulated proteins, encoded by mRNAs with 5′ terminal oligopyrimidine (TOP) tracts (**Figure 5D**, compare cyan and grey boxes). These data are consistent with the recent report that TOP mRNAs preferentially escape global suppression of translation by NSP1 (28).

Having established that our experiments identify known NSP1 binders and changes in total proteome, we also searched for possible novel interactors. Curiously, we found that WT NSP1 interacts with multiple components of the cellular anti-viral defense system (**Figure 5B**, red). The components we found include eukaryotic initiation factor 2 alpha (eIF2A, or eIF2S1) and eIF2A protein kinase R (PKR). PKR plays a protective role during viral infection: it is activated by doublestranded viral RNA, which leads to the phosphorylation of eIF2A and an inhibition of the synthesis of viral proteins (reviewed in (29)). We identified further interactions with a number of other antiviral components, including: NKRF (NF-kB -repressing factor), which mediates transcriptional repression of NK-kappa-B responsive genes (30); IFRD1 (interferon-related developmental regulator 1), which suppresses NF-kB activation (31); PDE12 (Phosphodiesterase 12), an enzyme that negatively regulates innate immunity (32); a number of helicases involved in IFN induction, including DDX21 and MOV10 (33,34); ILF3 (Interleukin enhancer-binding factor 3), required for translation of antiviral cytokine IFNB1 and a subset of INF-stimulated genes (10). These and additional interactions uncovered in our assay suggest that NSP1 may modulate anti-viral pathways via direct interactions with their components. Indeed, levels of some interactors, including DDX21, DDX1 and ILF3, were downregulated upon NSP1 expression, compared with the mock and KH164AA-expressing cells (**Figure 5C**). RK124AA mutant, that retains part of NSP1 repressive potential, showed fewer differences with WT NSP1 in respect with changing the levels of these binders (**Figure 5C**, WT/RK124AA). Curiously, we also observed that WT NSP1 downregulated other components of anti-viral defense pathway, including ADAR, involved in coronavirus genome editing (35); Bcl-2-associated transcription factor 1 (BCLAF1), that induces proinflammatory cytokines IL-6 and IL-8 (36); Annexin A1 (ANXA1), that upregulates cytoplasmic RNA sensor RIG-I and thereby stimulates IFNβ production (37).

NSP1 regulates both translation and stability, and RK124AA NSP1 mutant from SARS-CoV-1 (7) and SARS-CoV-2 (Biorxiv: https://doi.org/10.1101/2021.05.28.446204) was reported to be defective in mRNA destabilization. It has been speculated, that NSP1 might recruit a cellular nuclease in a manner dependent on intact R124 and K125 residues. We therefore looked for nucleases enriched among WT NSP1 interactors, compared with RK124AA mutant (**Figure 5B**, orange). Five nucleases have been identified: POP1, a component of ribonuclease P that generates mature tRNA by cleaving their 5′-ends (38); NOB1, an endonuclease required for processing of pre-rRNA precursor (39); ribosomal biogenesis protein LAS1L (40); DIS3, a component of the RNA exosome complex which possesses both 3′→5′ exoribonuclease and endonuclease activity (41); and 5′→3′ exoribonuclease XRN2 (42).

## Discussion

To date, the most prominent preventive approach to combat the impact of SARS-CoV-2 has revolved around the development of vaccines that target the viral spike protein. However, vaccination does not fully stop the propagation of disease, and new variants of SARS-CoV-2 may emerge which prove resistant to existing vaccines. This heightens the need for the development of drugs that target the core machinery of SARS-CoV-2 and can be used to treat infected individuals. NSP1 seems to be an ideal candidate as a target: it is conserved in beta-coronaviruses; it plays a crucial role in both downregulating the expression of host genes and promoting its own propagation; and it subdues the antiviral arsenal of infected cells. Assessing NSP1’s potential as a target will require the type of detailed mechanistic understanding of its functions that we present here.

The mechanism we describe offers an explanation for the way SARS-CoV-2 downregulates host mRNAs while concomitantly enhancing its own expression (**Figure 1 and 6**). The first step in this process is the early expression of viral protein NSP1. NSP1 blocks host translation by inserting its C-terminal domain into the mRNA entry tunnel on the ribosomal 40S subunit (12–14). Consistently, we detected interaction of WT NSP1, but not its nonfunctional mutant KH164AA, with ribosomal proteins and other components of the 43S pre-initiation complex (**Figure 5B**). Interestingly, a similar mechanism involving blocking of mRNA entry tunnel has been previously reported for other translational repressors, SERBP1 (43) and Stm1 (44).

**Figure 6.**
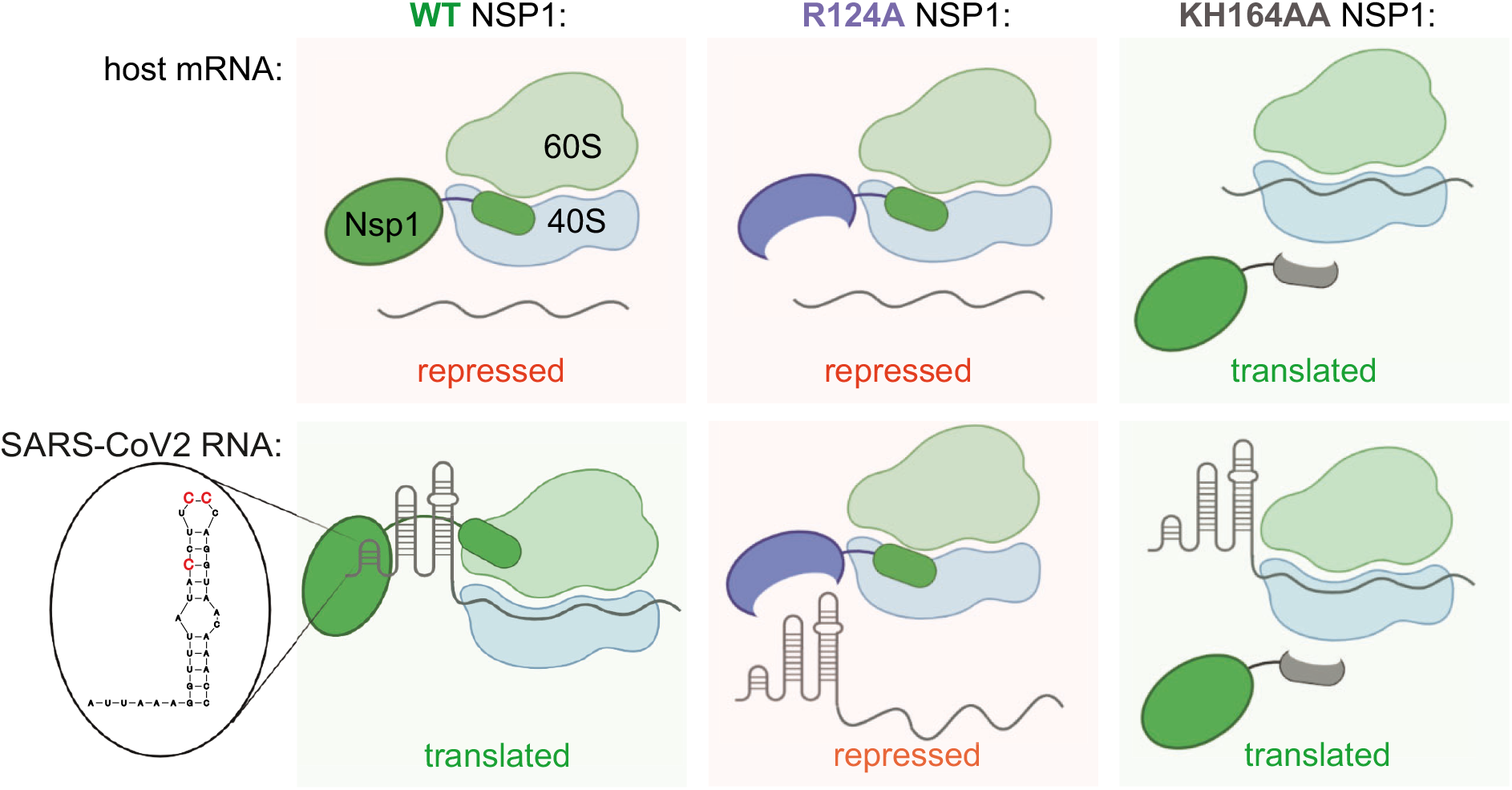
Speculative model for NSP1 function in repression of host mRNA and activation of SARS-CoV-2 expression. WT NSP1 protein represses translation of host mRNA via blocking the ribosome entry tunnel. SARS-CoV-2 genomic and subgenomic RNAs escape repression by NSP1 due to SL1 in their 5′UTR, interacting with NSP1. This way viral RNA highjacks host ribosomes without competing for limiting eIFs in infected cells. In particular, positions C15, C19 and C20 are required for alleviation of NSP1 silencing. Positions K164H165 in NSP1 are required for interaction with ribosome, therefore KH164AA mutant of NSP1 does not repress either host or SARS-CoV-2 mRNA. Position R124 is required for interaction with SL1, therefore NSP1 R124A mutant represses both host and viral mRNA.

Moreover, NSP1 interacts with the components of anti-viral machinery in cells (**Figure 5B**) in a way that suggests that it may directly hijack anti-viral pathways, beyond repressing the translation of host proteins by binding to the ribosome. In other cases, viral RNA is typically sensed by RNA helicases in infected cells, resulting in the activation of the transcription factors ATF2/c-Jun, IRF3/IRF7, and NF-kB (45). These in turn induce the production of cytokines, including members of the IFN family, which go on bind to their cognate receptors and trigger a second wave of signaling. These two waves upregulate genes that inhibit viral replication. By interacting with components of these pathways, NSP1 may prevent the full induction of IFNs as another mechanism that facilitates viral propagation. Indeed, we observed downregulation of some of these factors upon NSP1 expression (**Figure 5C**). Our NSP1 interactome data, including two mutants of NSP1 with fully or partially disrupted functionality, provide an important resource for future exploratory studies on SARS-CoV-2.

Previous findings that SARS-CoV-2 disrupts the production of cellular proteins left unanswered questions. To propagate, viruses must synthesize their own proteins, and this depends on the very machinery that appeared to be suppressed. Our work presents a mechanism by which SARS-CoV-2 ensures the translation of its own RNA, which depends on NSP1. We show that the stem loop SL1 within the viral leader sequence is both necessary and sufficient to escape NSP1-mediated repression (**Figure 2A**); this confirms findings by Banerjee et al. (16) and Tidu et al. (15).

Interestingly, Schubert et al. (13) failed to detect viral evasion in their reporter assays and suggested that virus may employ a different strategy. They proposed that viral transcripts might have a higher translation efficiency to begin with, which would give them a kinetic advantage in translation over cellular transcripts. Our data suggest a more likely explanation for the discrepancy in the two models. The viral leader used in Schubert’s study lacks the first 72 nt (according to the methods section of the manuscript), and therefore does not contain the SL1 required for viral evasion.

Interaction studies conducted with SARS-CoV-1 and SARS-CoV-2 NSP1 suggested that it can be bound by SL1 (9,15), Biorxiv: https://doi.org/10.1101/2020.09.18.302901). This likely causes NSP1 to be expelled from the mRNA entry tunnel. Curiously, Mendez et al. (Biorxiv: https://doi.org/10.1101/2021.05.28.446204) detected interaction of NSP1 with both viral and cellular reporter mRNAs, while earlier study of Tanaka et al. on SARS-CoV-1 (9) reports that viral leader is required for such interaction. Further structural studies will be required to resolve this discrepancy and uncover the specific mechanism behind the structural rearrangements of NSP1 on the ribosome that potentially occur upon its binding to SL1. Importantly, our analysis pinpoints specific residues within SL1 (three cytosine residues at the positions 15, 19 and 20) and NSP1 (R124) which are required for viral evasion and are likely involved in SL1/NSP1 interactions.

Our experiments revealed a dose-dependent response of viral reporters to NSP1. Specifically, at high doses of NSP1, viral reporters simply escaped silencing, but at low amounts, NSP1 actually stimulated their expression. We speculate that the enhancement of viral translation is caused by the global repression of host translation by NSP1. This generates a pool of translation factors and ribosomes that can now be coopted by the virus. This principle is known; it has been established for other viruses which carry internal ribosome entry sites and make use of a similar hijacking mechanism (reviewed in (46)). But this effect has not been previously reported for SARS-CoV-2 NSP1. The reason probably lies with the structure of prior studies. In some cases they have not compared expression of viral reporters in the presence and in the absence of NSP1 (16). Other studies have observed a similar expression of viral reporters under both conditions (Biorxiv: https://doi.org/10.1101/2020.09.18.302901, https://doi.org/10.1101/2021.05.28.446204), or detected a weaker repression of viral reporters, compared with non-viral reporters (15). The results of Shi et al. (Biorxiv: https://doi.org/10.1101/2020.09.18.302901) and Mendez et al. (Biorxiv: https://doi.org/10.1101/2021.05.28.446204) are consistent with our reporter assays in the presence of high levels of NSP1, and are easiest explained by high amounts of NSP1-encoding plasmid used in these studies. The most likely explanation for the effects observed by Tidu et al. (15) is that this work was performed in rabbit reticulocyte lysates, which are typically treated with nucleases to eliminate endogenous mRNAs. This system therefore does not recapitulate a possible competition for a limited number of translation factors, which is characteristic for translation *in vivo*.

An intriguing question remains regarding the extent to which NSP1 upregulates the expression of viral RNAs in the context of actual viral infection. Future *in vivo* experiments with viruses carrying mutant forms of NSP1 will be required to address this question.

NSP1’s fundamental role in viral infections make it a highly interesting potential target for drugs. The understanding we have gained of the mechanisms underlying its functions suggest three potential points of attack. The most obvious place to interfere is the site of the protein that interacts with the ribosome and blocks the mRNA entry tunnel. This is the defect observed in the KH164AA NSP1 mutant, which fails to interact with ribosome and is nonfunctional. Targeting K164 and H165 with small molecules therefore appear to be a promising strategy that would disrupt the pathogenicity of SARS-CoV-2. Another weak point that could be exploited is the mechanism which viral molecules use to evade NSP1 silencing: the structure that permits NSP1 to interact with the viral leader. There are two possible targets: the regions in either NSP1 or SL1 that permit and are required for this interaction. Our finding that the R124A mutant, but not WT NSP1, effectively represses the viral reporter points to a vulnerable spot on NSP1 that could be targeted by therapies. On the SL1 side, finding compounds that target three crucial cytosines (C15, C19 and C20), alone or in combination, might hold a great potential for the development of novel SARS-CoV-2 therapies.

## ACCESSION NUMBERS

The mass spectrometry proteomics data have been deposited to the ProteomeXchange Consortium via the PRIDE partner repository with the dataset identifier PXD024480.

## ACKNOWLEDGEMENT

Experiments were performed by L.B. (reporter assays Fig 1A, cloning, western blotting Fig 1A, 2B), M.C. (reporter assays Fig1B, 2A, 3 and 4, cloning, preparation of BioID samples), K.L. (cloning of BioID constructs), N. Z. (cloning of CoV-2-SL1 reporters), and D.K. (cloning of CoV-2-RL and RL reporters). O.S. and A.S. performed MS of BioID samples. N.v.K. performed statistical and exploratory data analysis for MS data. M.C. conceptualized and supervised the work, and wrote the paper, with the feedback from all authors. We thank Russ Hodge for the comments of the manuscript. Biorender.com was used in figure generation.

